# Robust metabolomics data normalization across scales and experimental designs

**DOI:** 10.1101/2025.09.30.679445

**Authors:** Matthijs Vynck, Pablo Vangeenderhuysen, Ellen De Paepe, Tim Nawrot, Vera Plekhova, Lynn Vanhaecke

## Abstract

Metabolomics studies employing liquid chromatography-mass spectrometry are affected by signal drift and batch effects, introducing technical variance that impedes biological knowledge discovery. Quality control (QC) sample-based normalization strategies are widely implemented but remain vulnerable to outliers, thereby reducing normalization performance. We introduce rLOESS, rGAM and tGAM, three robust normalization methods that improve resistance to outliers by downweighting or accommodating them. Leveraging additive models, the rGAM and tGAM methods allow flexible non-linear modeling, differential sample weighting, and data-driven QC representativeness evaluation. Implementations of these methods are gathered in the Metanorm R package, integrating robust normalization with visualization for performance verification, while supporting efficient parallel processing. In *in silico* and/or experimental datasets, the robust methods, relative to several popular existing strategies, improved replicate concordance, and reduced drift and batch effects. The robust methods, with improved recovery of the underlying signal demonstrated in simulation, produced distinct differential abundance results, highlighting the impact of normalization on downstream statistical inference. Overall, tGAM-based normalization suggested the best performance across scenarios and is proposed as default choice. Metanorm is versatile, supporting normalization in metabolomics studies across scales and experimental setups.

## Introduction

Liquid chromatography coupled to mass spectrometry (LC-MS) is a widely established analytical technique in metabolomics including lipidomics, and has supported significant discoveries across the life sciences and beyond^1–5^. As the technique has matured, its application has expanded to encompass increasingly larger biological sample series, nowadays frequently spanning hundreds if not thousands of samples^6,7^.

Analyzing such extensive sample series presents important analytical challenges. Indeed, extended analytical runs often require, e.g., intermediate instrument maintenance or consumable replacements. In addition, longer sample collection and analysis timeframes, often spanning weeks or months, introduce variability arising from factors such as multiple operators, shifts in sample handling, and environmental changes. Collectively, these factors exacerbate batch effects (e.g., column changes), and signal attenuation or enhancement, often referred to as drift, from both chromatographic and mass spectrometric origin, such as column degradation or fouling, as well as gradual changes in, e.g., ionization efficiency or mass calibration^8,9^. Consequently, these issues become more pronounced as sample series sizes increase.

Addressing this variability has become a critical concern in the metabolomics community, underscored by the establishment of the Metabolomics Quality Assurance and Quality Control Consortium (mQACC) in 2018^10^, which emphasizes the development of effective strategies to mitigate these challenges. While several strategies have been proposed, ranging from the use of isotopically labeled internal standards^11^ to broadly applicable statistical methods such as ComBat^12^, the most widely accepted approach today is the implementation of quality control (QC) samples to characterize and correct for batch effects and drift^9,13,14^. QC samples are typically generated by pooling aliquots from all, or in large sample series representative subsets of, biological samples^15^. Repeated analysis of these pooled QC samples allows for systematic monitoring and correction of batch effects and drift. Such QC sample-based corrections have become fundamental in supporting data reliability of large-scale LC-MS studies^9,10,14^.

Several QC sample-based normalization strategies have been proposed, including QC-RLSC, introduced by Dunn et al.^13^, employing locally estimated scatterplot smoothing (LOESS) for within-batch drift correction at the single metabolic feature level. This approach was later refined to a cubic spline-based method (QC-RSC) by Kirwan et al.^16^, reducing computational demands. More recently, methods such as MetaboQC^17^, SERFF^18^, and TIGER^19^ employed polynomial regression and LOESS, random forest regression, and ensemble learning strategies, respectively. A comprehensive listing and description are beyond this work’s scope.

Despite these advances, several challenges remain. In this work, we demonstrate that the presence of occasional, disproportionately high or low metabolite intensities (outliers) can deteriorate the performance of existing normalization methods. Outliers can occur infrequently, i.e., for a given (subset of) metabolite(s) (analyte outliers), or for all measurements in a sample (sample outliers); their treatment in this work does not differ. We subsequently show that such outliers go on to reduce statistical power in downstream analyses, such as differential metabolite abundance testing, likely contributing to numerous false negatives but also causing likely false positives. We introduce three robust approaches, built upon cross-validated robust LOESS, additive models (AMs), and generalized AMs (GAMs). We capitalize on (G)AMs’ ability for flexible non-linear modeling, for hypothesis testing and for observation weighting, refinements that enhance performance through a data-driven joint or selective usage of QC and biological samples for normalization. Given common pitfalls in benchmarking normalization strategies, such as using a reduction of the relative standard deviation of QC samples or an increased number of differentially abundant metabolites as a proxy for good normalization^20^, we propose and implement alternatives. To support application, we integrate both existing and the here introduced robust normalization methods into the Metanorm R package. Our implementations further capitalize on the problem’s pleasingly parallel nature, resulting in efficient high-throughput normalization. In addition, the package provides complementary visualization options that support normalization performance verification.

## Experimental Section

### Current normalization strategies

For benchmarking we compared the three robust normalization approaches with the current QC-RLSC and QC-RSC strategies. Given that no metabolomics community-wide accepted reference implementations of these approaches exist^14^, they were implemented based on details provided in the original manuscripts^13,16^, insofar these details were available. Specifically, for QC-RLSC we used the loess function (stats package) with degree set to two and implementing a leave-one-out cross-validation (LOOCV) for span selection, with a span in the range 𝟑/𝒏 ≤ 𝜶 ≤ 𝟏, 𝒏 being the number of QC sample injections. The optimal span is selected as that span minimizing the mean-squared error of left-out observations. As spans below approximately 0.075 occasionally yield model fitting problems on experimental datasets, the minimal span taken is the highest value in the set {𝒏/𝟑, 𝟎. 𝟎𝟕𝟓}. An alternative, faster implementation using generalized cross-validation (GCV) was also implemented, relying on the loess.as function in the fANCOVA package. For QC-RSC, following Kirwan et al.^16^, we applied the smooth.spline function (stats package) of the R language, whose implementation allows for smoothness selection using either LOOCV or GCV, with default restrictions on the smoothness parameter.

For the evaluation on experimental datasets, the QC-RLSC and QC-RSC approaches, as well as SERFF^18^ and TIGER^19^ were included in the comparative analyses together with our three robust methods. SERFF was used through its Shiny graphical user interface provided by Fan et al.^18^, at https://slfan.shinyapps.io/ShinySERRF/, whereas TIGER was available as an R package (Supplementary Table 1).

### Three robust normalization methods

To increase robustness against outliers, the first normalization method uses a robust locally estimated scattersplot smoothing (LOESS) approach. Robustness is achieved by (iteratively) downweighing outlying observations, with weights determined by Tukey’s biweight function (as implemented in the loess.as function in the R fANCOVA package). The optimal span is determined using either GCV or LOOCV.

Further normalization flexibility, while retaining robustness against outliers, is achieved by adopting one of two distinct (generalized) additive model (GAM) formulations.

The first, which we named the robust GAM (rGAM) method, achieves robustness in a similar manner to the rLOESS approach. For a given compound, an initial standard GAM fit is obtained for the model

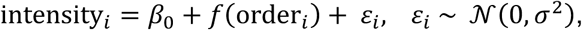

with *i* = 1, …, *n* the (QC) injection order index, *β*_0_ an intercept, *f* a smooth function, and 𝜀_*i*_ a Gaussian error term. Next, observation weights are calculated based on the initial model fit’s residuals, using Tukey’s biweight function,

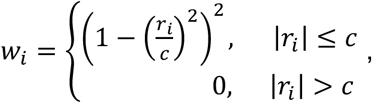

with *r_i_* the fitted model’s rescaled residuals, i.e. the model’s residuals divided by a scaled median absolute deviation (MAD) of the residuals (MAD ∗ 1.4826), and 𝑐 = 4.6821, a constant^21^. The model is then re-fitted with these observation weights *w_i_,*

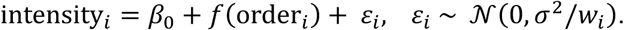

The second strategy, which we named the tGAM method, achieves robustness more directly by adopting a scaled-𝑡 distribution for the outcome,

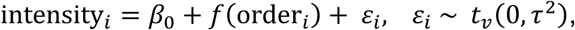

with *t_v_* a scaled-𝑡 distribution with 𝑣 degrees of freedom. The use of a scaled-𝑡 distribution rather than a Gaussian distribution reduces the impact of outliers on the estimated intensity for a given injection order^22^.

Of note, while traditionally only QC samples are used^13,16^, smooth function estimation can be based on QC samples alone, biological samples alone, or their combination.

Optionally, for the additive models, observation weights can be supplied to assign more importance to any given observation during model fitting, e.g., assigning higher weights to QCs than samples so that the former exert a higher influence on the estimation of the smooth function.

### Additive model refinements for dealing with batch effects

The additive model-based methods were further extended to properly handle batch effects. To achieve this, a batch covariate was added to the GAM model:

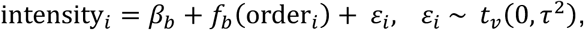

where 𝑏 = 1, … , 𝐵 denotes the 𝐵 batches, i.e., the smooth function *f_b_*, as well as the intercept *β_b_* is allowed to differ by batch. Note that the scaled-𝑡 error term is replaced by a Gaussian error term, depending on the chosen model (tGAM *vs.* rGAM). Due to the computational complexity that arises when many batches are involved, and because the residual variance between batches may differ, a ‘batchwise’ option was implemented: in this setting, an additive model is fitted per batch data subset, reducing model complexity and allowing for batch-specific residual variances,

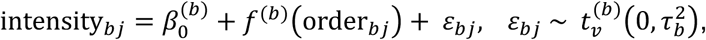

with 𝑗 = 1, … *n*_’_ a batch-specific index, *n*_’_ the number of runs in batch 𝑏, β_0_(b) and *f*(b) a batch-specific intercept and smooth function respectively, and 𝜀*_bj_* a batch-specific error term (again, potentially replaced by a Gaussian error term depending on the chosen model).

### Additive model refinements for verifying whether QCs are representative of biological samples

A final optional adjustment enables an evaluation of (within-batch) differences in intensities between biological samples and QCs. This is achieved by adding a ‘type’ covariate to the GAM model specification, indicating whether a run concerns a QC or a biological sample, resulting in separate smooth functions for biological samples and QCs:

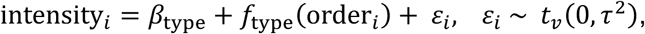

where *β_type_* and *f_type_* denote type-specific intercepts and smooth functions, respectively. Significant differences, i.e., a *p*-value lower than a user-specified cut-off, between QC- and sample-based intercepts and/or smooth functions, indicating unrepresentative QC samples, are determined using generalized likelihood ratio tests^23^, and will then result in a model re-fit and normalization using biological samples only.

### Implementation

The three robust methods rLOESS, rGAM and tGAM, along with the existing methods QC-RLSC and QC-RSC were implemented in the Metanorm R package (available from https://github.com/UGent-LIMET/Metanorm). Users can select a method of choice using the ‘model’ argument, with options ‘QC-RLSC’, ‘QC-RSC’, ‘rLOESS’, ‘rGAM’ or ‘tGAM’. To reduce over- or underfitting, smoothness parameter estimation for the rLOESS, QC-RLSC and QC-RSC approaches are available with either LOOCV or GCV schemes (‘cv’ argument). All GAMs are fitted using restricted maximum likelihood^24^ and thin plate regression splines (R mgcv package (Supplementary Table 1)), with the basis dimension (‘k’ argument) user-specified. Refinements can be combined in a single call of the normalization algorithm, i.e., observation weights (‘weights’ argument), the modeling of batch effects (‘batch’ argument), and QC versus biological sample modelling by using the ‘QCcheck’ argument for enabling comparison, and the ‘QCcheckp’ argument for setting the significance threshold.

### Experimental data

Three experimental datasets were selected to investigate the impact and performance of the different normalization strategies. The first is the publicly available metabolomics BioHEART study dataset^25,26^. The BioHEART dataset comprises 1359 human plasma sample runs (of which 1002 unique biological samples) across 15 batches, quantifying the abundance of 53 targeted metabolites using an Agilent 1260 Infinity LC coupled to a QTRAP 5500 Sciex MS instrument. Data were preprocessed using Sciex’ MultiQuant. Further details are provided by^25^. The occurrence of differentially abundant metabolites (DAMs) in this dataset was investigated for individuals with (n=390) or without (n=612) hypertension^25,26^.

Secondly, a targeted metabolomics dataset from the ENVIRONAGE cohort was used^27^. A total of 322 children aged 4 to 12 years were included, of whom 48% (n=156) were boys. This dataset spans 393 human urine sample runs (of which 322 unique biological samples) across 6 batches, acquired using a Dionex Ultimate 3000 XRS UHPLC system (Thermo Fisher Scientific) coupled to a Q-Exactive MS instrument (Thermo Fisher Scientific). Further analytical details are provided by De Paepe et al.^28^. The resulting data were preprocessed using TARDIS^29^, and metabolites with more than 10% missing values were removed. The final dataset used for assessing normalization performance comprised 269 targeted metabolites. DAMs in relation to biological sex were investigated^30^. Of note, batches in this dataset did not always start with a QC sample; as QC-RLSC and QC-RSC cannot extrapolate beyond QCs within batches, samples analyzed before the first QC in each batch could not be included for both methods.

Thirdly, an untargeted metabolomics dataset from the FAME cohort was utilized^31^. The FAME dataset encompasses 618 human saliva sample runs (of which 486 unique biological samples), extracting 8913 metabolic features, acquired with a Vanquish Horizon ultra high performance LC system linked to an Orbitrap Exploris 120 MS (Thermo Fisher Scientific). Data preprocessing was performed using Thermo Fisher Scientific’s Compound Discoverer 3.3 software. The occurrence of DAMs related to a healthy weight (337 individuals) *vs.* overweight/obesity (99 individuals) was evaluated. Forty-four cases had no data for weight status at the time of analysis and were excluded from the DAM analysis.

All intensities were log-transformed (base *e*) prior to normalization.

### Comparing performance using simulated data

Drift was simulated *in silico* by sampling two hypothetical QC sample intensity values per 10 biological samples, for a total of 1000 samples. This resulted in 84 QC sample intensities. Intensities were taken from a sine function, offset by a randomly chosen value of 20 to avoid negative intensities, after which Gaussian noise (standard deviation 0.3) was added:

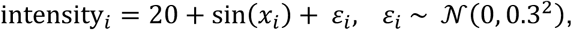

with 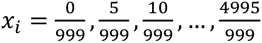, chosen to represent part of a full sine period. The dataset was then split into two equal parts, to obtain two distinct drift patterns (Supplementary Fig. S1). Next, zero, one, two or four outliers were added, with the following characteristics: (i) outlier locations were selected randomly but could not occur in the first five or last five observations, and (ii) outliers were created by subtracting between 20 and 30% (randomly selected within that range), subtracting between 20 and 100%, or subtracting 100% or adding 200% of the simulated intensity value. Outlier parameters were feature-specific, i.e., differed between each simulation run, allowing to evaluate their impact across sizes and locations in the analysis sequence.

In each of 𝑘 = 200 simulation runs, encompassing simulation of a single feature, and for a given number of outliers, QC-RLSC, QC-RSC, rLOESS, rGAM and tGAM models were used for data normalization, using both one and two QC samples per 10 biological samples. Performance was then assessed by calculating, for each method, the squared difference between the predicted and expected drift pattern at all QC indices in each simulation run. Finally, the median across all runs was calculated, yielding an estimated median squared prediction error (MSPE) for each of the five methods, having used one or both QC samples, and zero to four outliers, i.e., a total of 40 MSPE estimates.

### Comparing performance using experimental data

Significant differences between groups of interest (see section Datasets) after normalization were compared between methods. First, significant DAMs were determined using two-sided Wilcoxon rank sum tests, with *p*-values < 0.05 taken to indicate significance. A comparatively equal number of DAMs, and especially a notable overlap in DAMs was taken to demonstrate similar normalization performance. In contrast, much larger or smaller numbers of DAMs, or low overlap was taken to demonstrate a deviating normalization performance, which was subsequently further investigated at the compound level.

Additionally, fine-grained comparability was assessed by calculating the pairwise Spearman correlations of the methods’ *p*-values. Similarity was then quantified as the pairwise Euclidean distance of one minus the correlation vectors of any two normalization methods. These were visualized in a heatmap, with rows and columns arranged according to a hierarchical clustering with complete linkage to support an assessment of comparative or divergent normalization performance across metabolites, as manifested in terms of biological signal discovery.

In the BioHEART study, a subset of biological samples was run in replicate in subsequent batches. As normalization was performed batchwise, no between-batch information leakage occurred, thus allowing to use the consistency of observed intensities of these replicate runs as a benchmark for appropriate normalization. Consistency was assessed by first calculating the difference between the two observed intensities (rescaled by their mean intensity) in each pair of replicate runs, for all compounds, then obtaining the MAD of these observed differences per normalization approach. A lower MAD then indicated better correspondence of intensities across batches, suggesting superior normalization.

Finally, after removal of runs with missing data, two-dimensional principal component analysis (PCA) score plots were constructed to allow a qualitative evaluation of batch effects and outliers pre- and post-normalization.

### Computational time and compute details

All five methods implemented in the Metanorm package, TIGER, through its R package, and SERFF through its web interface, were evaluated for computational run times across scenarios. These scenarios involved both targeted (53 compounds, BioHEART) and untargeted (8913 features, FAME) experiments, large (1361 samples, BioHEART) and medium (618 samples, FAME) sample sets, and large (15 batches, BioHEART) and medium (9 batches, FAME) batch numbers. Computational time was determined for analyses using from 1 to 6 compute cores, to support an assessment of the potential time gains due to the parallel computing approach implemented in the Metanorm package. Note that this was not possible for SERFF as it was run through its web interface. All methods were run with default arguments and parameter settings, except for the rGAM and tGAM methods that were run with the QCcheck argument active. To prevent central processing unit throttling affecting the time required for the sequential analyses of different methods, a two-minute break between any two analyses was ascertained. Computational time required was expressed as the number of normalized compounds per minute.

All analyses were performed using a 10-core Apple M1 Pro processor and 16GB RAM. The analyses were performed using R version 4.4.1^32^. Package details are provided as supplementary information (Supplementary Table S1).

## Results

### Three outlier-robust metabolomics normalization methods

This work introduces three normalization methods allowing outlier-robust signal intensity drift and batch effect correction. The first and simplest method modifies the QC-RLSC method^13^ by downweighting outliers, thereby reducing their influence on the signal intensity drift fit (Fig. 1a). This robust locally estimated scatterplot smoothing (rLOESS) model further incorporates generalized cross-validation (GCV) as a replacement for the traditionally used leave-one-out cross-validation (LOOCV)^33^. The other two robust methods employ (generalized) additive models (GAMs) for signal drift and batch effect correction. The first GAM-based method, a robust GAM (rGAM) downweights outliers to minimize their impact on the signal drift fit, analogous to the rLOESS method (Fig. 1a). The second GAM-based method adopts a *t*-distribution (tGAM) rather than a Gaussian distribution for the intensities, enabling to handle outliers effectively, as the *t*-distribution’s heavier tails reduce the influence of these outlying intensities (Fig. 1a). The outlier-robustness of our three methods exceeds that of QC-RLSC, QC-RSC and SERFF, which showed inferior normalization performance in batches containing outliers, exemplified in Fig 1a (see below for additional results). Results from our evaluation on *in silico* generated data causally confirmed the superior normalization performance of our three robust methods, both in the absence and presence of outliers, but in particular in the latter situation. Moreover, the robust methods remained comparatively unaffected by the number of outliers and magnitude of deviation. (Fig. 1b, S2-S4).

**Figure 1:**
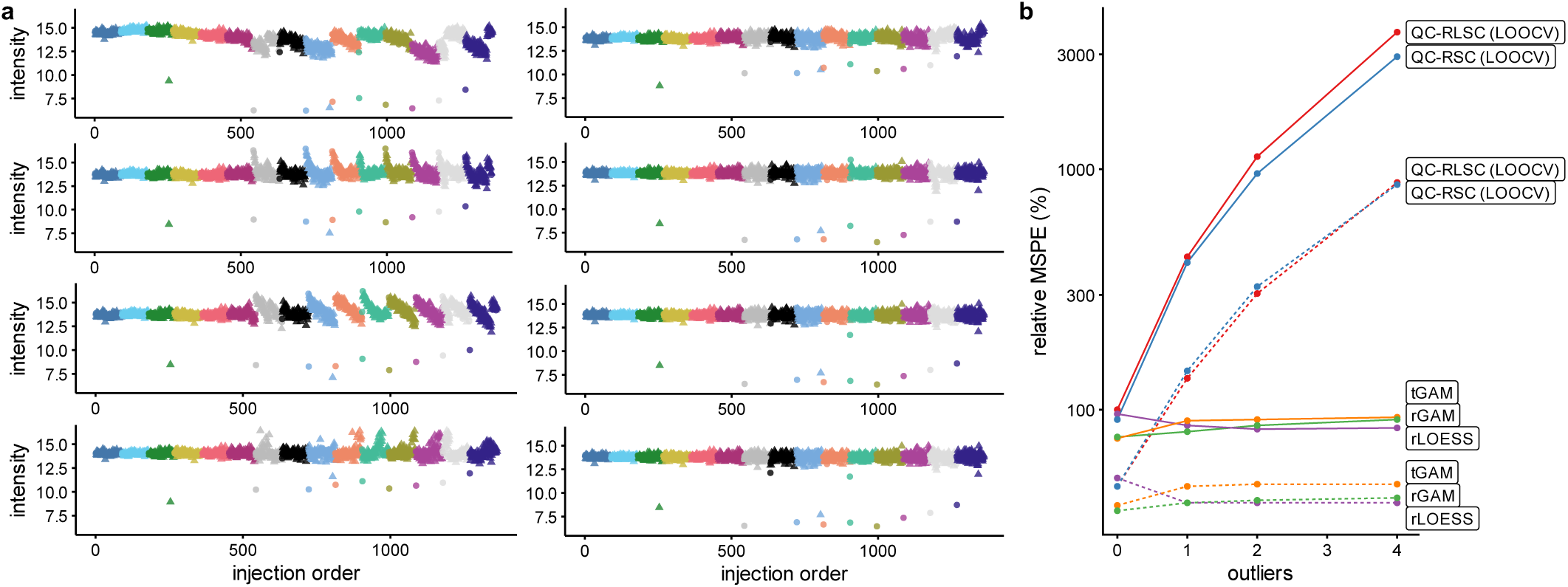
Susceptibility of normalization method to outliers and batch effects. (a) Unnormalized, QC-RLSC, QC-RSC, SERFF normalized (left column, top to bottom), and TIGER, rLOESS, rGAM or tGAM normalized (right column, top to bottom) for an example metabolite from the BioHEART study (see below for additional results). Dots indicate QCs, whereas triangles indicate samples; colors correspond to different batches. (b) Mean squared prediction error (MSPE (%), relative to QC-RLSC with zero outliers) for normalization methods as a function of the number of outliers for in silico generated data. Full line: 1 QC per 10 samples; dashed line: 2 QCs per 10 samples. [Note: two-column figure]

The rLOESS, rGAM and tGAM methods are implemented in the Metanorm R package (https://github.com/UGent-LIMET/Metanorm), alongside implementations of the QC-RLSC and QC-RSC methods.

### Flexible modelling boosts normalization performance: leveraging convoluted variance in biological samples, and data-driven normalization

While QC-sample-based normalization is widely established, biological samples themselves also carry information on signal drift patterns and batch effects. The tGAM and rGAM implementations in the Metanorm package enable differential weighting of QC and biological samples during normalization. By assigning greater weight to QC samples, technical variance predominantly drives estimation of the normalization smooth, while batch- and signal-related information from biological samples is still incorporated (Fig. 2a-2c).

**Figure 2:**
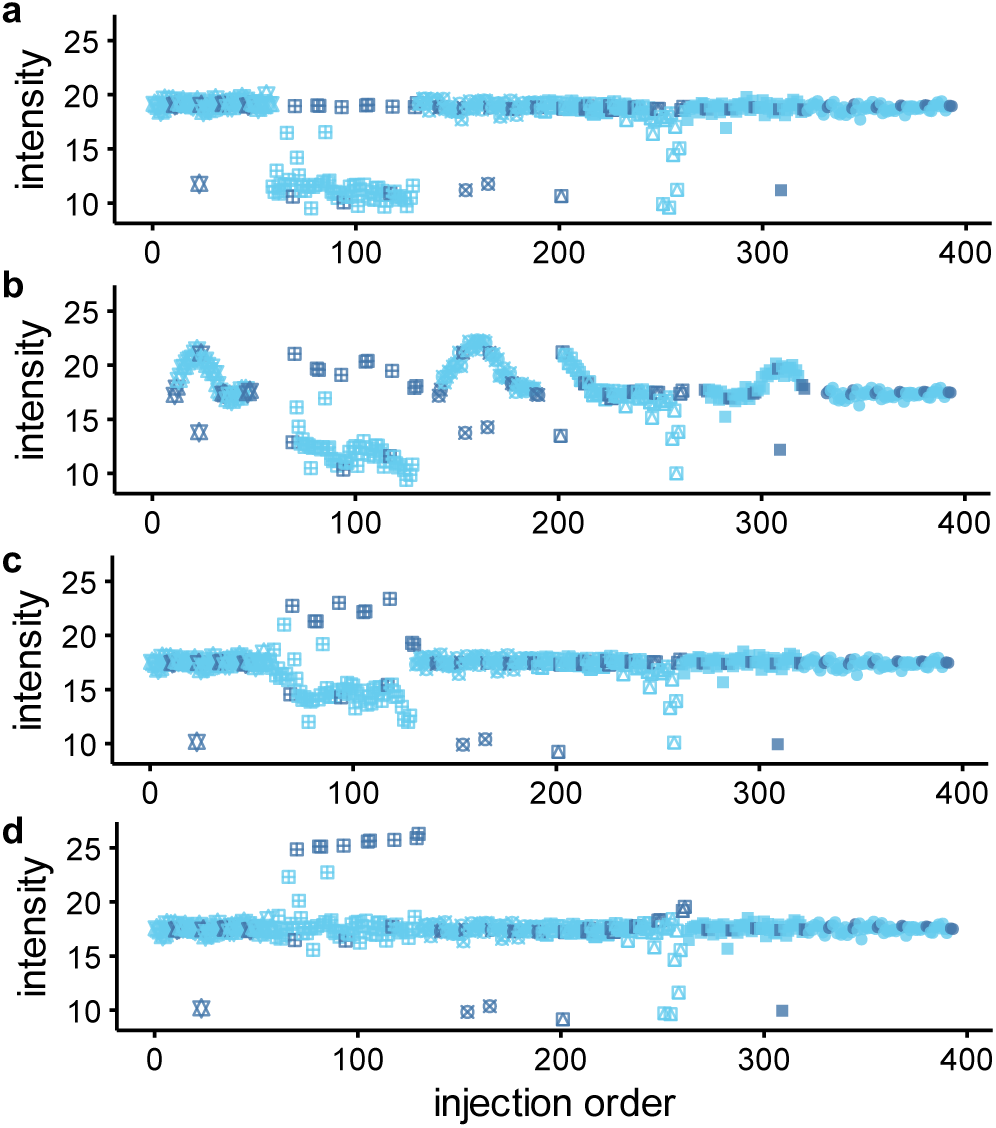
Impact of outliers and data-driven QC- versus biological sample-based normalization for the metabolite Cyclo(Leucylprolyl) in the ENVIRONAGE urine analysis. Dark blue symbols indicate QCs, light blue symbols indicate biological samples; symbol types correspond to different batches. (a) Unnormalized intensities. (b) QC-RLSC normalized intensities – affected by outliers in the QC samples. (c) tGAM normalized with a five-to-one weight ratio of QC vs. biological samples but without QC versus biological sample check – affected by a difference in QC and biological samples. (d) tGAM normalized intensities with a five-to-one weight ratio of QCs versus biological samples and with QC versus biological sample check. [Note: single-column figure]

Further, the Metanorm package supports visual assessments of normalization performance. Diagnostic plots of pre- and post-normalization compound abundances can be automatically generated (Fig. 2). These plots highlight whether normalization successfully reduces technical variance across batches or, conversely, whether it fails to do so. An issue recurring across datasets was that QC-based normalization did not sufficiently remove technical variance or, in some cases, even introduced additional variance in biological samples due to differences between QC and biological samples (Fig. 2a-2c, S5). Given the infeasibility of verifying, and potentially adjusting, QC-based normalization for every single compound and batch, especially in untargeted studies, the Metanorm package includes an optional statistical test (QCcheck argument) to assess the suitability of QC-based normalization at the single compound (and batch) level. This test, compatible with both GAM-based methods, models signal drift separately for QCs and biological samples within each batch. If significant differences between the two are detected, the rGAM/tGAM normalization reverts to sample-based rather than combined sample and QC-based normalization (Fig. 2d). The stringency of this decision is controlled using the QCcheckp parameter. Together, these functionalities and diagnostic tools allow to evaluate and finetune normalization, while maximally extracting signal drift and batch effect information.

### Validation on experimental data suggests robust normalization reduces false positive and false negative findings, and improves reproducibility

Experimental results confirmed the improved normalization performance of our three robust methods previously observed in the *in silico* datasets.

The analysis of the BioHEART dataset indicated that the tGAM, rGAM and rLOESS methods resulted in the detection of largely the same DAMs (92% joint DAM findings, Fig. 3b-3c). SERFF, however, showed strong disagreement with the robust methods (29-41% joint DAM findings; Fig. 3b-3c). Investigation of pre-and post-normalization intensity-versus-order plots for these discrepant findings supported proper normalization of the robust methods, which were little affected by outliers (Fig. 3a). In contrast, SERFF was impacted by few outliers, often showing increased variance for those biological samples in proximity to such outlying observations (Fig. 3a). This is supported by the PCA score plots, which show a strong reduction in batch effects compared to non-normalized data for all normalization methods, but also small residual batch effects for SERFF, which are not seen for the tGAM method. (Fig. 3d). Support for outliers as the root cause for this sub-par performance was provided. Indeed, manual identification and removal of outliers improved SERFF’s normalization performance, with reduced variance observed in the proximity of previous outlying intensities (Supplementary Fig. S6). Not only SERFF resulted in an increased number of DAMs: notably, the unnormalized, raw data showed the highest number of DAMs, followed by TIGER and the QC-R(L)SC approaches (Fig. 3b). In all cases, discrepancies in both significant and non-significant findings between the robust and other methods could be linked to remaining drift or batch effects (Fig. 3a, Supplementary Fig. S7-S8). Further supporting the enhanced normalization performance of the robust methods, the normalized intensities of between-batch sample replicates showed the lowest between-replicate normalized differences, i.e., the highest correspondence, for the robust methods (49-52% reduction in between-replicate normalized differences, compared to unnormalized data), followed by SERFF (41% reduction), TIGER (40% reduction), then QC-RLSC and QC-RSC (18-22% reduction; Fig. 4).

**Figure 3:**
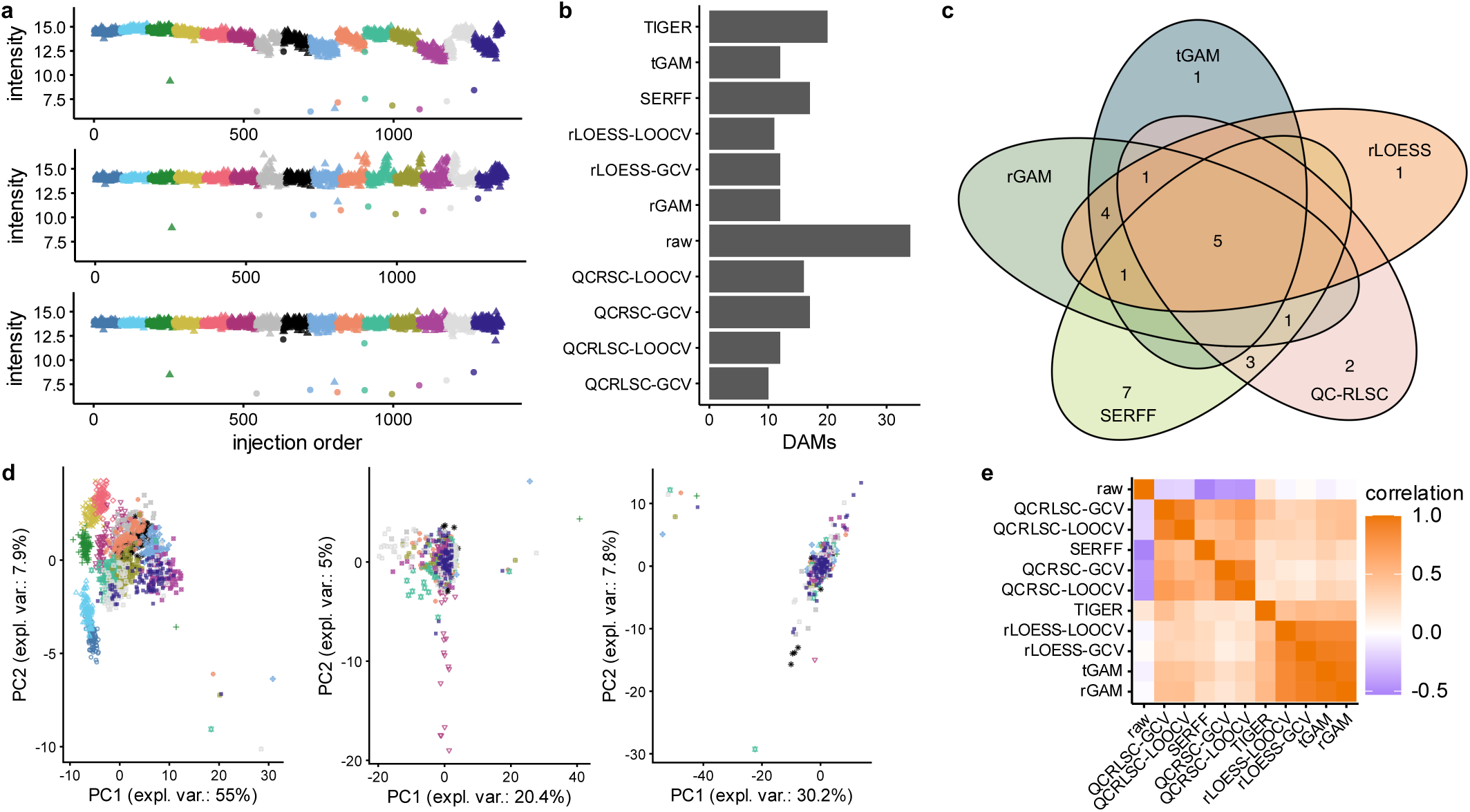
Impact of normalization methods on the BioHEART data. (a) Methionine intensities before (top), after SERFF (middle), and after tGAM (bottom) normalization (Figure S7 for all methods). Dots indicate QCs, whereas triangles indicate biological samples; colors correspond to different batches. (b) Number of significantly differentially abundant metabolites (DAMs) by normalization method. (c) Venn diagram of DAMs for the three robust methods, SERFF and QC-RLSC. (d) Principal component score plots before (left), after SERFF (middle), and after tGAM (right) normalization. Symbol types and colors correspond to different batches. (e) Heatmap of correlations of p-values as obtained by Wilcoxon tests, per normalization method; arranged according to hierarchical clustering. [Note: two-column figure]

**Figure 4:**
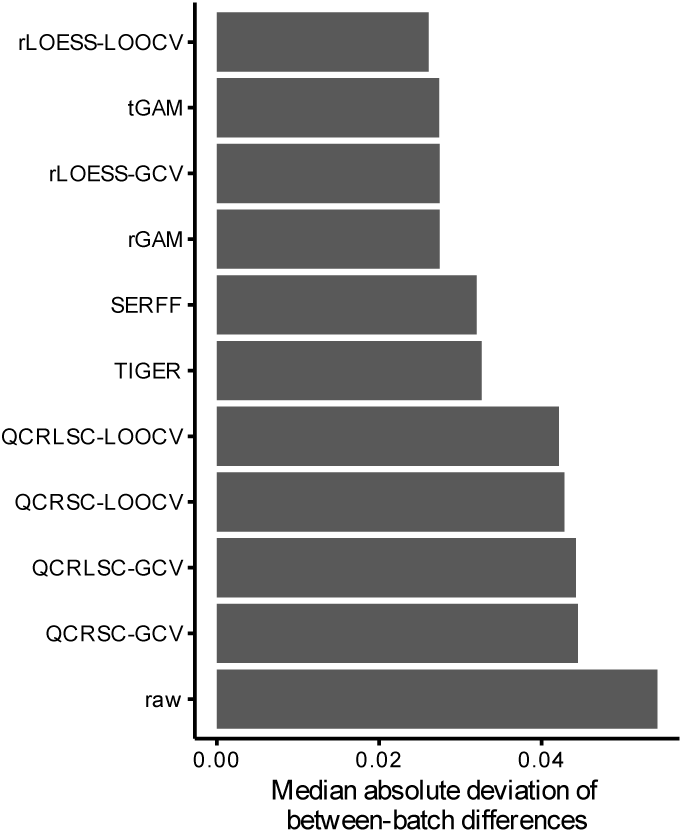
Median absolute deviation of scaled between-batch differences of compound intensities in non-QC samples for which two replicate runs were performed in the BioHEART study. Lower median absolute deviation indicates a better cross-batch agreement, and suggests an improved removal of technical variance. [Note: one-column figure]

For the ENVIRONAGE dataset, similar albeit less pronounced patterns were observed (Fig. 5), likely explained by the comparatively smaller batch and drift patterns, and the fewer and less severe outliers seen when compared to the BioHEART dataset (Fig. 5a, 5d). Specifically, the robust methods were in reasonable agreement (74-88% DAM overlap, Fig. 5b-5c), whereas QC-RLSC and, to a lesser extent, SERFF and TIGER differed more strongly from the robust methods (47-51%, 68-70% and 64-67% DAM overlap, respectively, Fig. 5b-5c). Differences in significant DAMs were mostly due to relatively small differences in *p*-values, illustrated by comparatively high cross-method correlations (Fig. 5e), though occasional larger differences arose due to fitting of QC-RLSC to non-representative QC observations, and due to SERFF’s sensitivity to outliers (Fig. 5a and Supplementary Fig. S9).

**Figure 5:**
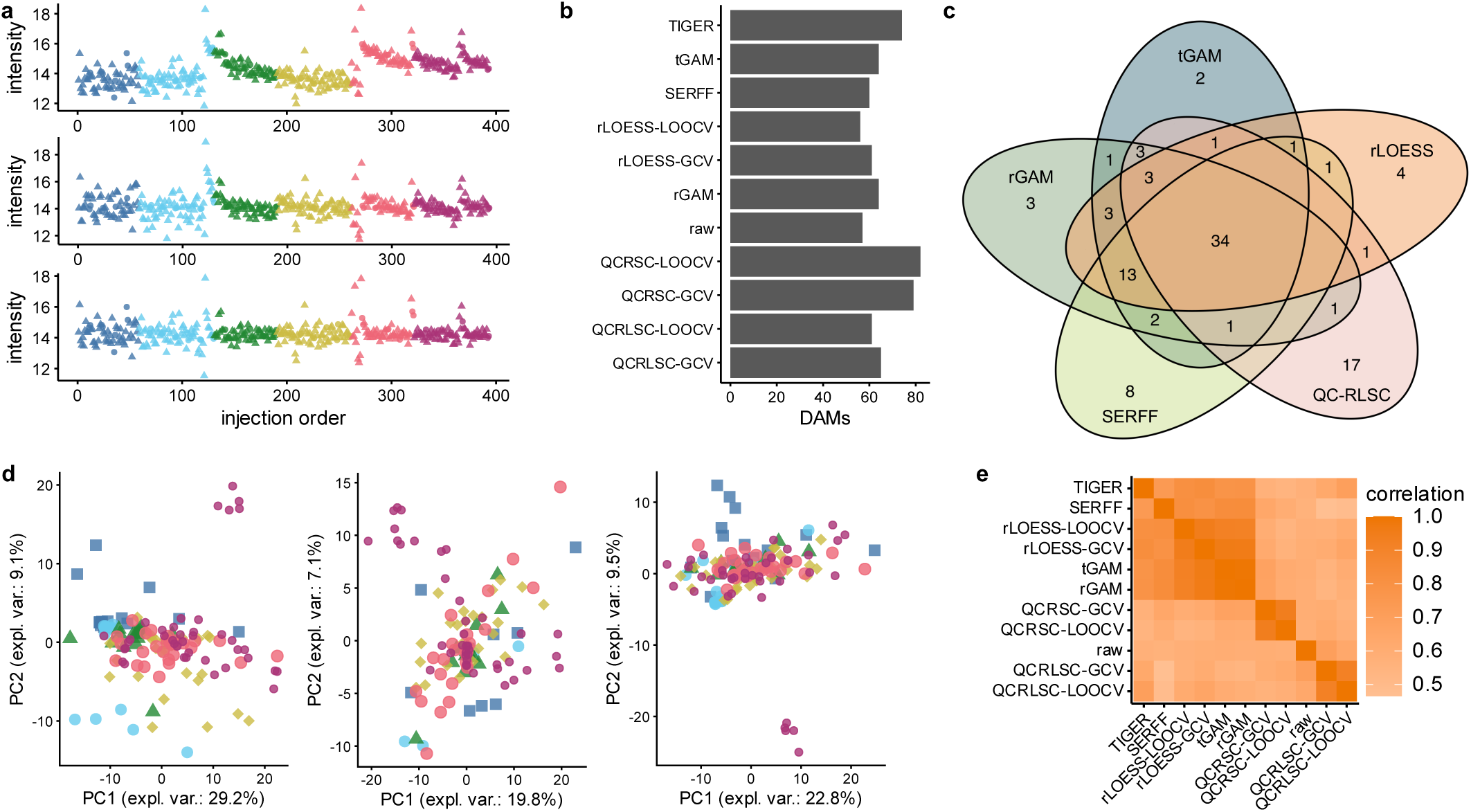
Impact of normalization methods on the ENVIRONAGE data. (a) Phenylglycine intensities before (top), after SERFF (middle), and after tGAM (bottom) normalization. Dots indicate QCs, whereas triangles indicate biological samples; colors correspond to different batches. (b) Number of significantly differentially abundant metabolites (DAMs) by normalization method. (c) Venn diagram of DAMs for the three robust methods, SERFF and QC-RLSC. (d) Principal component score plots before (left), after SERFF (middle), and after tGAM (right) normalization. Symbol types correspond to different batches. (e) Heatmap of correlations of p-values as obtained by Wilcoxon tests, per normalization method; methods arranged according to hierarchical clustering. [Note: two-column figure]

For the FAME cohort, again, good agreement between the robust methods was observed (79-87% DAM overlap), whereas QC-RLSC showed weaker agreement (55-59% DAM overlap; Supplementary Fig. S10). Disagreements between QC-RLSC and the robust methods could often be traced to differences in QC *vs.* biological sample intensities (Supplementary Fig. S9). No results for SERFF could be obtained, likely due to the size of the dataset exceeding the server’s capacity.

Across datasets, qualitative comparisons of pre- and post-normalization intensity-versus-order plots between the three robust methods suggested that the rLOESS method was more likely to show signs of overfitting. This was evidenced by occasional more wiggly smooth functions when compared to the rGAM and tGAM methods (Supplementary Fig. S11). Differences between the rGAM and tGAM methods were tentatively traced to the tGAM method at times being more robust to (groups of) outliers than the two-step rGAM method (Supplementary Fig. S12).

Spanning all datasets and methods, the choice of cross-validation scheme had comparatively little impact, the number of DAMs and the strength of evidence for differential abundance between either GCV or LOOCV being highly comparable within normalization methods (Fig. 3b, 3e, 5b, 5e, Supplementary Fig. S6).

### Robust and flexible normalization is computationally expensive, but offset by a parallel computing implementation

An assessment of the computation requirements showed stark differences: QC-RSC was fastest overall, with the rLOESS and QC-RLSC with GCV also delivering fast. Using six-core processing, all three methods normalized the BioHEART targeted data (53 compounds) in under 10 seconds (Fig. 6a) and the FAME untargeted data (8913 compounds) in under 7 minutes (Fig. 6b).

**Figure 6:**
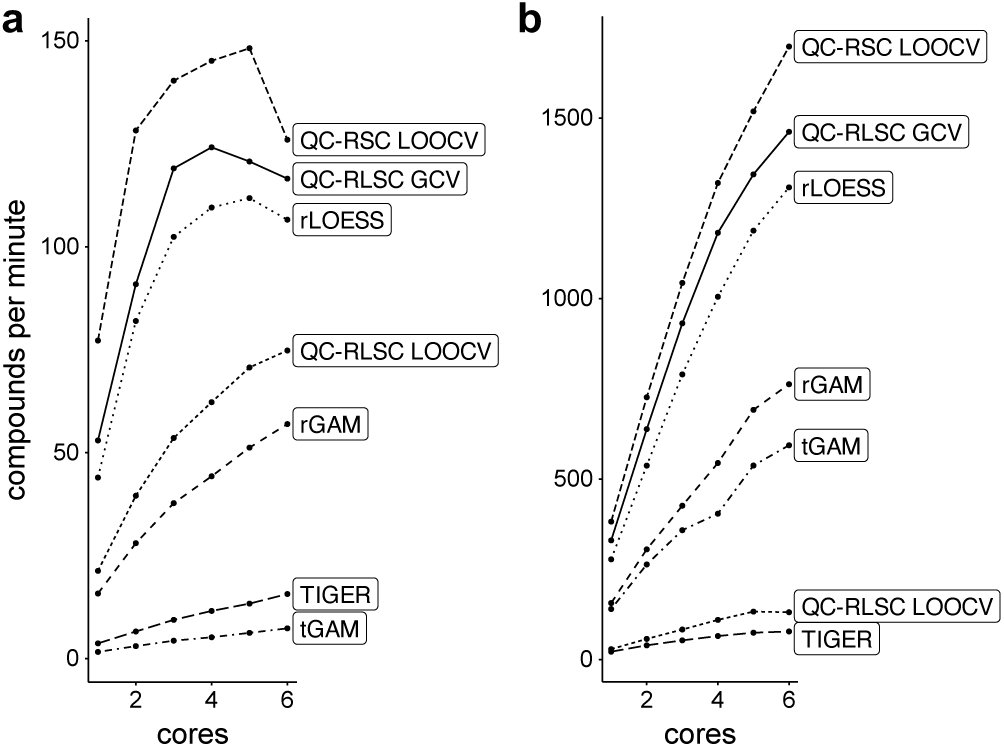
Time required for compound normalization by method. (a) BioHEART targeted data. (b) FAME untargeted data. [Note: single-column figure]

The more complex methods, TIGER and the GAM-based approaches, took notably longer, taking approximately 18 (rGAM), 64 (TIGER) and 136 (tGAM) seconds for the BioHEART dataset (Fig. 6a), and approximately 11 (rGAM), 15 (tGAM) or 115 (TIGER) minutes for the FAME dataset (Fig. 6b). The difference in ordering noted between both datasets could be explained by differences in the number of runs (1361 *vs.* 618), and/or the number of batches (15 *vs.* 9). SERFF, only available through its web server, took 147 seconds for the BioHEART dataset and failed to complete for the FAME dataset.

The computational overhead associated with parallel processing resulted in marginally slower normalization for the fastest methods in the BioHEART dataset, as evidenced by the tapering and then declining curves as more cores were used (Fig. 6a). This was not seen for the larger FAME dataset, where, across methods, more cores led to a steady decrease in computational time required. This suggests that, for datasets encompassing many compounds, an even higher number of cores than used here would further expedite data normalization (Fig. 6b).

## Discussion

The three normalization methods introduced here, rLOESS, rGAM and tGAM, are founded in well-studied statistical models^23,24^, and their implementation in the Metanorm R package facilitates robust, high-throughput, and efficient normalization. By integrating robust non-linear modelling with differential sample weighting and data-driven modelling into a single framework, along with visualization options, the Metanorm package supports reliable normalization of both small- and large-scale metabolomics data, as evidenced in both *in silico* and experimental data analyses, consistently across instruments, sample types, and laboratories. The implied reductions in false positives and false negatives observed in the experimental analyses, attributed to reduced susceptibility to outlying intensities, strengthen the reproducibility of downstream statistical analyses, unlocking more efficient biological knowledge discovery.

Overall, the tGAM method exhibited the strongest performance across the experimental datasets, whereas performance on simulated data was comparable between rLOESS, rGAM, and tGAM. A practical limitation of tGAM is its higher computational cost relative to rLOESS and rGAM; however, normalization remains computationally feasible, completing within minutes on current consumer-grade hardware even for datasets comprising thousands of metabolites and hundreds of samples. Taken together, and based on the evidence presented here, we therefore tentatively recommend the tGAM method as a general default choice. Comparison of multiple methods remains useful as method performance may be dataset-specific.

An alternative to the use of the outlier-robust normalization procedures introduced here could be outlier detection and removal prior to normalization. Such outlier removal would need to be automated, as manual identification of outliers is both subjective and labor-intensive for typical metabolomics datasets often constituting hundreds to thousands of features. While automated outlier detection is non-trivial^34^, it is a potential alternative to the procedures introduced in this work. In particular, multivariate outlier detection may improve the current feature-level treatment of outliers. It is important to note that an observation that reduces normalization performance may still be correct and thus warrant exclusion or downweighting during normalization, but not necessarily during downstream statistical inference.

Besides the methods’ increased robustness to outliers, we have shown that, while being widely applied^10,14^, relying solely on QC sample-based normalization poses threats and misses opportunities. First, QC-based normalization assumes that QC samples ubiquitously and accurately mirror the technical and biological variance in study samples. Data originating from different instruments and laboratories investigated in this work show that this is not a universal truth. Such differences may arise from, e.g., different pre-analytical treatment of QC *vs.* biological samples, repeated injection of the same QC sample, etc. When this assumption of QCs being representative of biological samples does not hold, QC-based normalization may not only underperform but may even reduce data quality by introducing artefacts. By adaptively testing and, when necessary, reverting to a sample-based normalization strategy, Metanorm’s GAM-based procedures protect against such underperformance and artefacts. This adaptability is particularly important for large-scale untargeted metabolomics studies, where the representativeness of QC samples is difficult to verify manually across thousands of features. Second, biological samples themselves contain exploitable information on drift and batch effects, not leveraged when using only QC samples for normalization. By including the larger number of biological samples compared to QC samples (often a 5-10-fold difference^14^), the available information on signal drift and batch effects is substantially increased, although present in biological samples as a convolution of technical and biological variance. The implementation of differential weighting in the tGAM and rGAM methods provides a compromise: by giving QC samples stronger influence while still incorporating the drift information embedded in biological data, a more precise normalization can be achieved. This is particularly useful to support normalization in scenarios where QC sample coverage is sparse or uneven, arising when, e.g., QC sample analysis has failed, or when QC samples are lacking altogether. Visualization tools further enhance quality control, allowing to inspect whether normalization has achieved its desirable effects. The pre- *vs.* post-normalization and PCA score plots readily generated by the Metanorm package support an efficient identification of subpar data quality or aberrant normalization, and can highlight metabolites, samples or batches that require further, detailed inspection. We emphasize that unexpected differences between QC samples and biological samples should receive due analyst scrutiny, as such differences may also reflect issues with the biological sample analysis. In addition, any differences between QC and biological samples may invalidate filtering on e.g., relative standard deviation (RSD) observed in QC samples, if such differences are due to issues with the QC samples, discussed next.

In terms of verifying appropriate normalization, traditionally, the workhorse metric is the RSD, also known as the coefficient of variation (CV)^13,16,18,19^, with substantially reduced RSDs after normalization being taken as a reflection of good performance. Nonetheless, the RSD has been critiqued for favoring overfitting^20^.

Indeed, some strategies, while having merit and reducing technical variance, can result in small RSDs due to fitting non-signal drift related random noise^35^. The same is true when no proper cross-validation is conducted. We avoided such evaluation metrics measures when assessing Metanorm as we believe such surrogate measures are ideally replaced by those measures with a causal relationship with normalization performance. One such approach broadly applied in the statistics and bioinformatics literature but, to our knowledge, underexplored for (QC-based) data normalization methods in metabolomics is through *in silico* simulation studies where the ground truth is known^36^. Such an approach was incorporated in this study but could for future studies be further enhanced to more comprehensively verify normalization performance. This could be achieved by better mimicking experimental metabolomics data, for example, by simulating from observed data using resampling strategies, while accounting for cross-metabolite correlations. Indeed, such a strategy would allow to also benchmark multivariate methods as SERFF and TIGER. In addition, benchmarking a broader set of normalization methods remains important future work. In benchmarking, care should be taken to assure neutrality, i.e., not potentially favor any of the benchmarked methods which can arise from, e.g., simulating from a fitted model from one of the benchmarked methods^37^. Other evaluation metrics should equally be interpreted with due care. For example, the extent to which biological signal can be extracted does not necessarily correlate with normalization performance (e.g., De Livera et al.^20^). As with RSDs, this is an indirect evaluation approach, and findings in this work suggest that not only false negatives but also false positives could arise due to subpar normalization, a finding that is also supported elsewhere^37^. An important nuance, for method benchmarking, p-value thresholding was used for method ranking and relative comparisons, not for formal hypothesis testing with controlled error rates. All normalization strategies were applied to the same feature set and evaluated under a single nominal threshold. Applying a multiple testing correction would alter the number of selected features but is unlikely to affect (i) the relative ordering of normalization strategies, or (ii) the overlap patterns that were the primary focus of our evaluation. Indeed, conclusions are based on consistency and overlap of selected features across normalization methods; the number of significant features should not be interpreted as a proxy of normalization quality as inappropriate normalization can induce false positives and false negatives. While not in the scope of this work, multiple testing correction is essential for downstream statistical inference. Last, we employed technical replicates for evaluating normalization performance. When these replicates are sufficiently spaced, e.g., analyzed in distinct batches, information leakage is prevented when using per-batch normalization methods. Replicate concordance is thus also expected to be an unbiased measure of normalization accuracy. We recommend inclusion of both QC and replicate samples to support data quality and normalization performance investigation.

Our methods focus on interpretable normalization at the metabolite level. Fairly recently, machine learning models for metabolomics data normalization have been proposed and tentatively shown to outperform existing approaches^18,19^. While these methods have notable strengths, such as their capability to model potential interactions between signal drift patterns across metabolites, they also suffer potential drawbacks. For example, these approaches were demonstrated to have good performance on large datasets (with up to thousands of samples, and several hundreds of metabolites), but their reliability for smaller datasets has so far not been broadly investigated. In contrast, our metabolite-level normalization methods perform equally well on targeted datasets investigating few metabolites and on untargeted datasets, irrespective of whether they encompass dozens or thousands of samples. Further enhancements of metabolite-level approaches, however, could be achieved by leveraging cross-metabolite information.

## Conclusion

Current approaches to normalization of metabolomics data can be sensitive to outliers in metabolite intensities, leading to subpar normalization, and eventually false negative and false positive discoveries. Three new methods, rLOESS, rGAM and tGAM, that are robust to such outliers were presented, with evaluations suggesting improved normalization performance. Additional flexibility, such as the ability to handle occasional discrepancies between biological samples and QCs in an automated way, can further enhance normalization performance. Critically, normalization method selection was shown to strongly affect the number of differentially abundant metabolites, with, consequently, potentially large effects for downstream mechanistic knowledge discovery. These new methods, together with commonly used alternatives, are implemented in the Metanorm R package, which exploits parallelization for efficient processing. Visualization of metabolite intensities before and after normalization remains important for assessing performance, and options supporting such performance assessments are included in the Metanorm R package.

## Supporting information

Supporting Information: Table of R packages and versions used (Table S1); supporting figures: an example simulated dataset (Figure S1), normalization performance on simulated data (Figures S2-S4), selected feature plots illustrating pre- and post-normalization feature intensities in a variety of settings (Figures S5-S9, S11-S12), and additional results of normalization performance on the FAME cohort data (Figure S10).

## Data availability

The targeted metabolomics data from the BioHEART cohort is available at https://github.com/SydneyBioX/BioHEART_metabolomics. The targeted metabolomics data from the ENVIRONAGE cohort is available at https://zenodo.org/records/14548033. The untargeted metabolomics data from the FAME study is available at the Metabolomics Workbench with project ID PR002287 (http://dx.doi.org/10.21228/M80Z5F), under study ID ST003697.

## Code availability

The Metanorm package is available at https://github.com/UGent-LIMET/Metanorm. The code used for the analyses and generating figures is available at https://github.com/UGent-LIMET/Metanorm_publication.

## Acknowledgements

The authors thank the Laboratory of Integrative Metabolomics’ administrative and technical personnel for their continuous support of the lab’s research activities.

## Author contributions

MV conceptualized and developed the ideas, implemented the methods, performed data analysis, created figures, wrote the initial draft and edited the draft paper. PV contributed to the development of the ideas and reviewed and edited the draft paper. EDP and VP contributed to the generation of the data, provided feedback on the software, and reviewed and edited the draft paper. TN contributed to the generation of the data, and reviewed and edited the draft paper. LV acquired funding, contributed to the development of the ideas, supervised the project, and reviewed and edited the draft paper.

