## Supporting Information for "Robust metabolomics data normalization across scales and experimental designs"

### Table of Contents

Table S1 R packages and versions used *p.* S2
Figure S1 Example of simulated dataset *p.* S3
Figure S2 Examples of normalization and data-generating curves, no outliers *p.* S3
Figure S3 Examples of normalization and data-generating curves, with outliers *p.* S4
Figure S4 Impact of outlier severity on the relative mean squared prediction error (MSPE) *p.* S5
Figure S5 Normalized intensities for Cyclo(leucylprolyl) in the ENVIRONAGE dataset *p.* S5
Figure S6 Effect of removing two outlying observations on SERFF's normalization results *p.* S6
Figure S7 Methionine intensities before and after normalization with all methods in the BioHEART dataset *p. S7*
Figure S8 Six cases of aberrant normalization in the BioHEART dataset *p.* S8
Figure S9 Four examples of aberrant normalization in the ENVIRONAGE and BioHEART datasets *p.* S9
Figure S10 Impact of normalization methods on the FAME dataset *p.* S10
Figure S11 Comparison of rLOESS and tGAM, metabolite ID86 in the FAME dataset *p.* S10
Figure S12 Comparison of rGAM and tGAM, metabolite ID101 in the ENVIRONAGE dataset *p.* S11

*Table S2: R packages and versions used*

| Package | Package version |
|---|---|
| changepoint | 2.3 |
| dplyr | 1.1.4 |
| fANCOVA | 0.6-1 |
| futile.logger | 1.4.3 |
| ggplot2 | 3.5.2 |
| mgcv | 1.9-3 |
| nlme | 3.1-168 |
| parallel | 4.4.1 |
| pbapply | 1.7-2 |
| RandomForest | 4.7-1.2 |
| readxl | 1.4.5 |
| reshape2 | 1.4.4 |
| splines | 4.4.1 |
| TIGERr | 1.0.0 |
| VennDiagram | 1.7.3 |

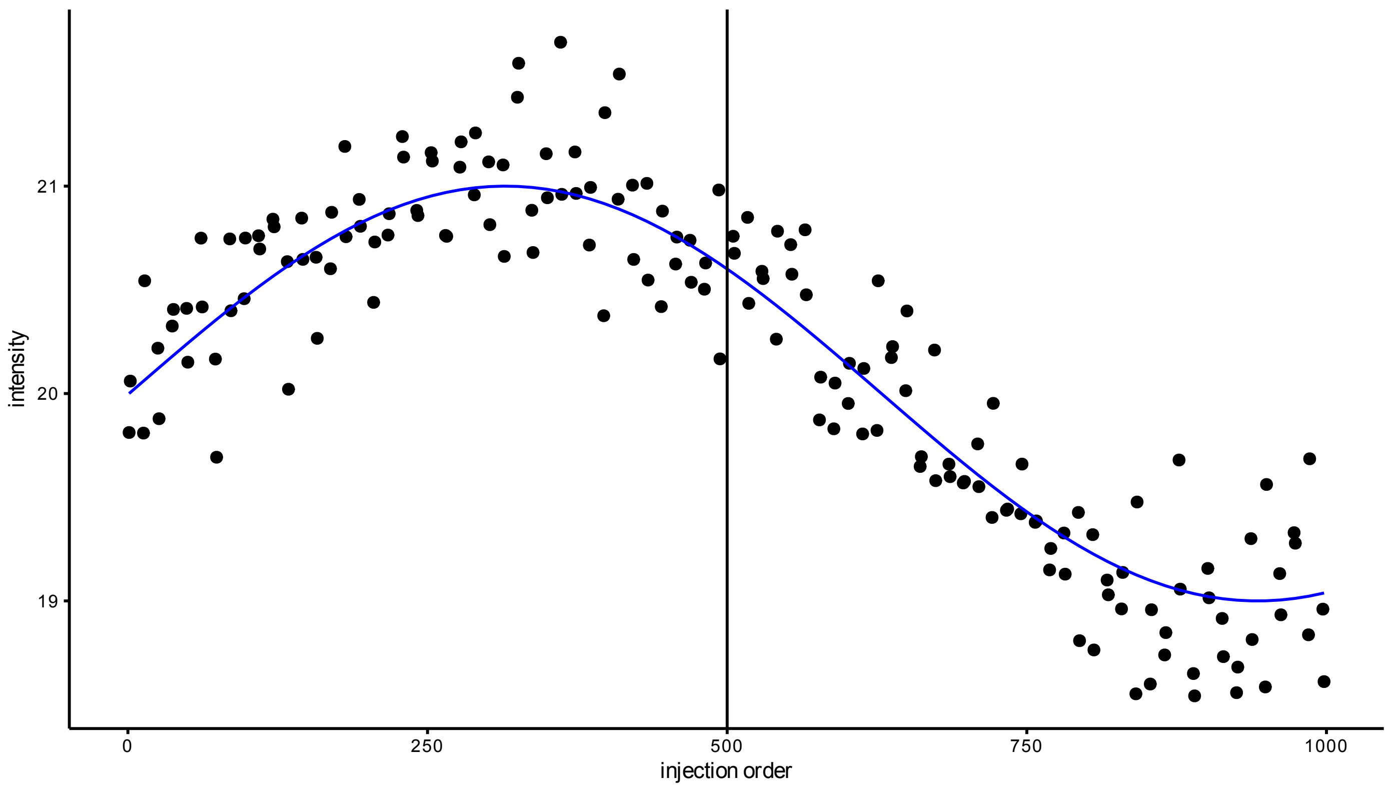


*Figure S1: Example simulated dataset. The black vertical line separates the two known drift functions used for data simulation (blue line). Black dots represent the simulated observations, obtained by adding Gaussian noise to the known drift function. Zero, one, two or four outliers are then added to assess robustness of data normalization approaches to outliers.*

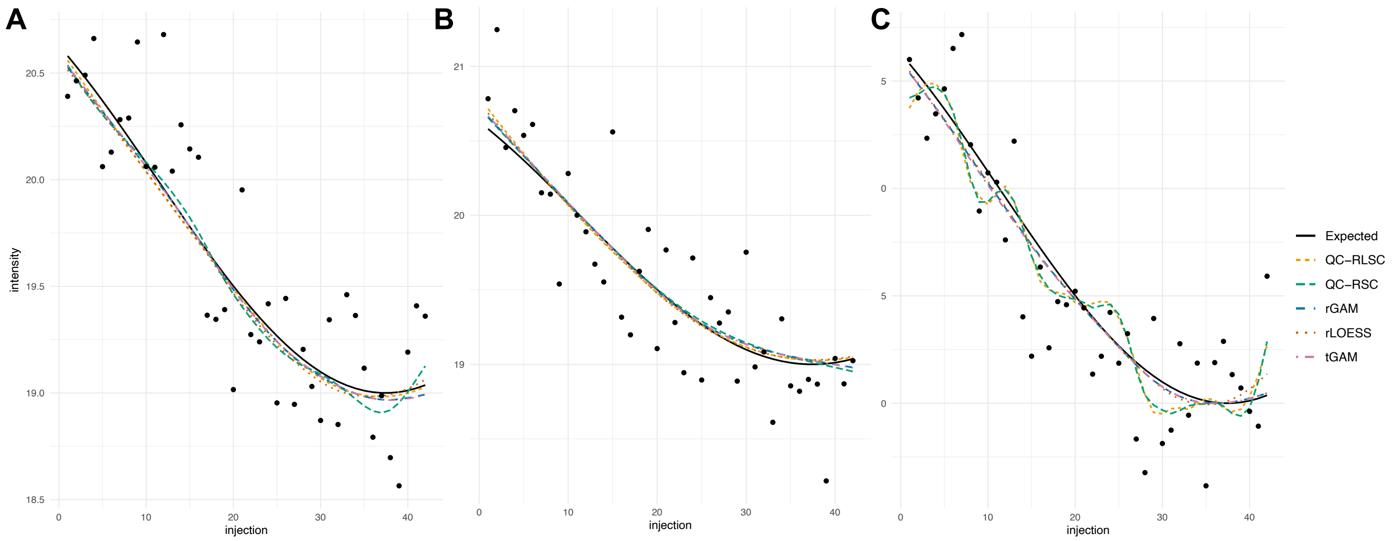


*Figure S2: Examples of normalization curves (colored) and data-generating curve (black), with simulated observations (points). Without outliers (see Figure S3 for examples with outliers), normalization curves are in the majority of cases in good agreement with the data-generating curve and with one another (panels A, B), however, occasionally, QC-RLSC and QC-RSC and to a lesser extent rLOESS show comparatively high disagreement with the data-generating curve, fitting with their higher RMSE (panel C).*

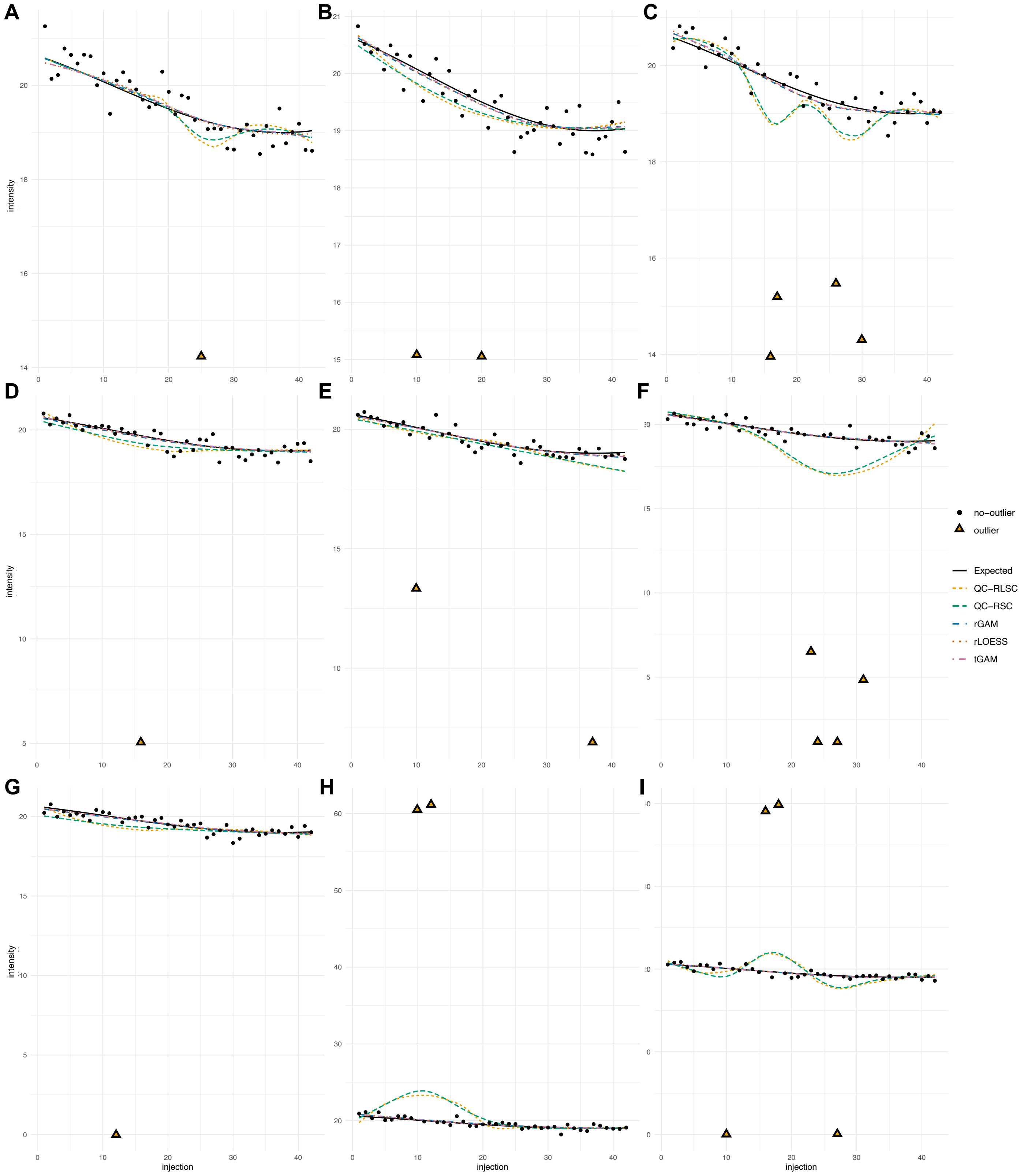


*Figure S3: Examples of normalization curves (colored) and data-generating curve (black), with simulated observations (points). Outliers (triangles) were introduced by (A-C) decreasing the observations by 20-30%, (D-F) decreasing the observations by 20-100%, (G-I) decreasing the observations by 100% or increasing them by 200%. Different numbers of outliers were added: 1 (A, D, G), 2 (B, E, H) or 4 (C, F, I). Location and severity of the outliers affect how much the curves obtained by non-robust normalization approaches deviate from the data-generating curve, whereas outliers do not heavily affect the robust methods. Occurrence of outliers in proximity to one another generally has a larger influence (e.g., panels C, F, H, I) then when they are more spaced (e.g., panels B, E).*

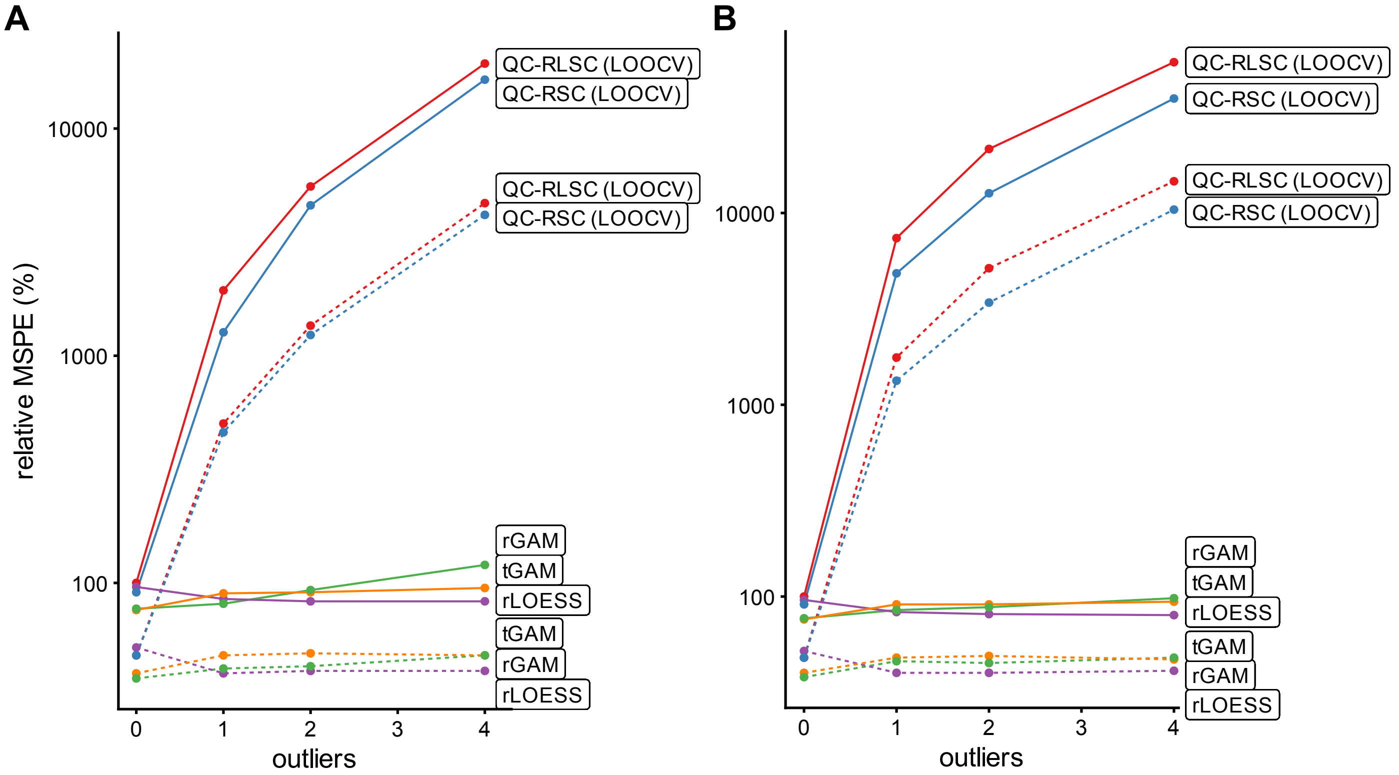


*Figure S4: Impact of outlier severity on the relative mean squared prediction error (MSPE) of the normalization approaches. (A) With observations 20-100% lower than originally observed. (B) With observations 100% lower or 200% higher than originally observed. Stronger outliers result in higher MSPE for QC-RLSC and QC-RSC (i.e., MSPE in panel A is lower than in panel B for 1, 2 or 4 outliers), whereas the robust methods remain largely unaffected.*

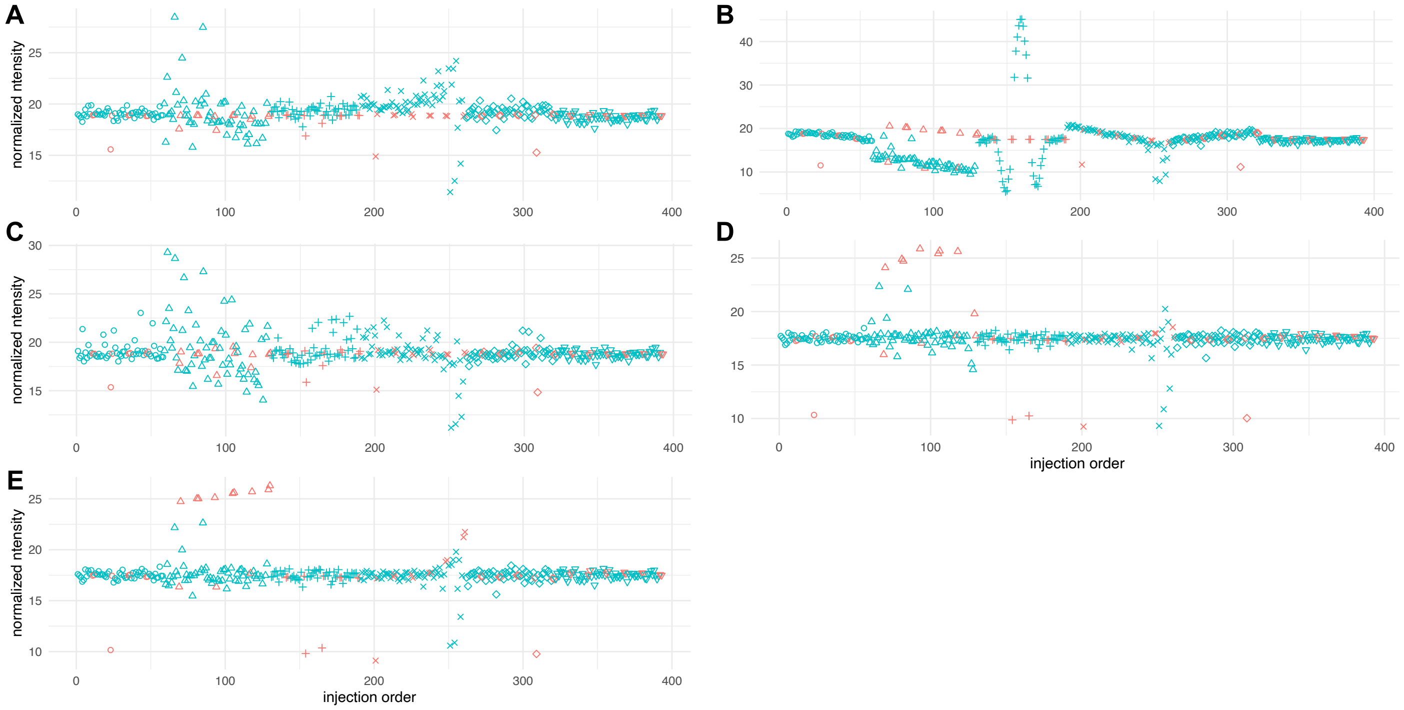


*Figure S5: Normalized intensities for Cyclo(Leucylprolyl) in the ENVIRONAGE urine analysis; results of QC-RLSC and tGAM with different weighting strategies is shown in Figure 2 in the main text. Blue symbols indicate samples, coral-red symbols indicate QC samples; symbol types correspond to different batches. (a) TIGER normalized, suggesting adequate correction of batch effects, (b) QC-RSC normalized, suffering from QC-biological sample differences, and moreover showing comparatively large sample deviations in the third batch, fitting with the methods' higher RMSE (c) SERFF normalized, suggesting adequate correction of batch effects but showing larger variance in the second batch, (d) rLOESS normalized, suggesting adequate removal of batch effects, (e) rGAM normalized (no weighting), suggesting adequate removal of batch effects.*

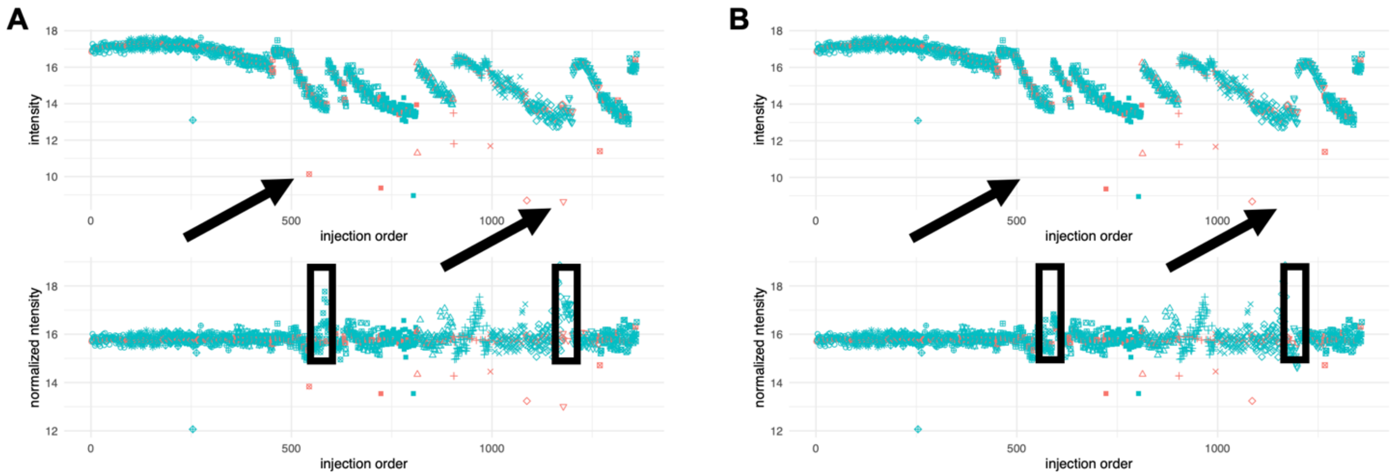


*Figure S6: Effect of removing two outlying observations on SERFF's normalization results. Blue symbols represent biological samples, coral-red symbols represent QC samples. Different symbols represent batches. (A) Original data before (top panel) and after (bottom panel) SERFF normalization. (B) Adjusted data, the two observations indicated by the black arrows removed, before (top panel) and after (bottom panel) SERFF normalization. Note that part of the above-average intensities in the normalized data have disappeared.*

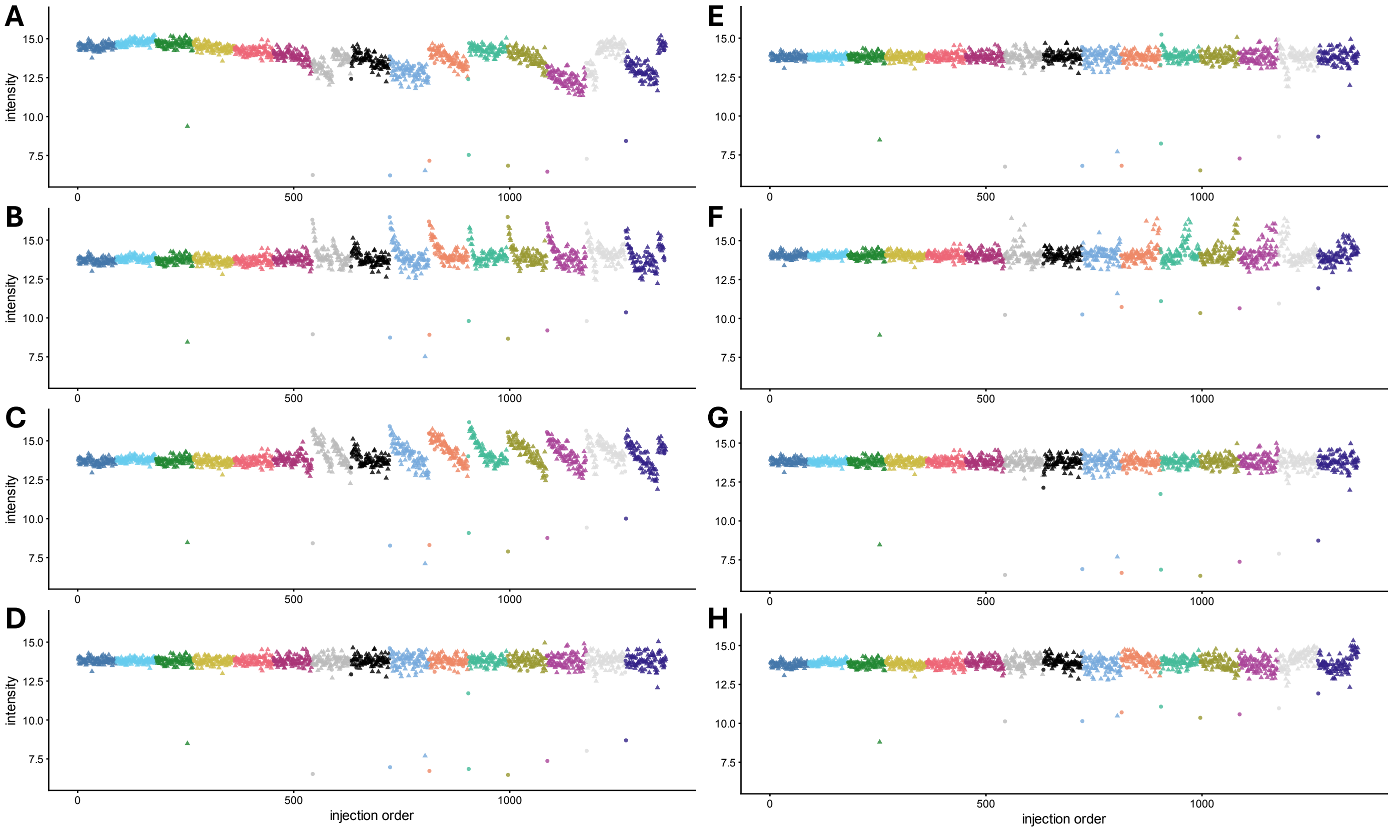


*Figure S7: Methionine intensities in the BioHEART dataset before normalization (A); and after normalization using different methods (B-H); colors represent different batches.*
*From top to bottom in alphabetical order, qualitative interpretation:*
*(B) QC-RLSC suggesting remaining within-batch drift*
*(C) QC-RSC suggesting remaining within-batch drift*
*(D) rLOESS suggesting comparatively low remaining within-batch drift*
*(E) rGAM suggesting comparatively low remaining within-batch drift*
*(F) SERFF suggesting remaining within-batch drift*
*(G) tGAM suggesting comparatively low remaining within-batch drift*
*(H) TIGER suggesting intermediate remaining within-batch drift*

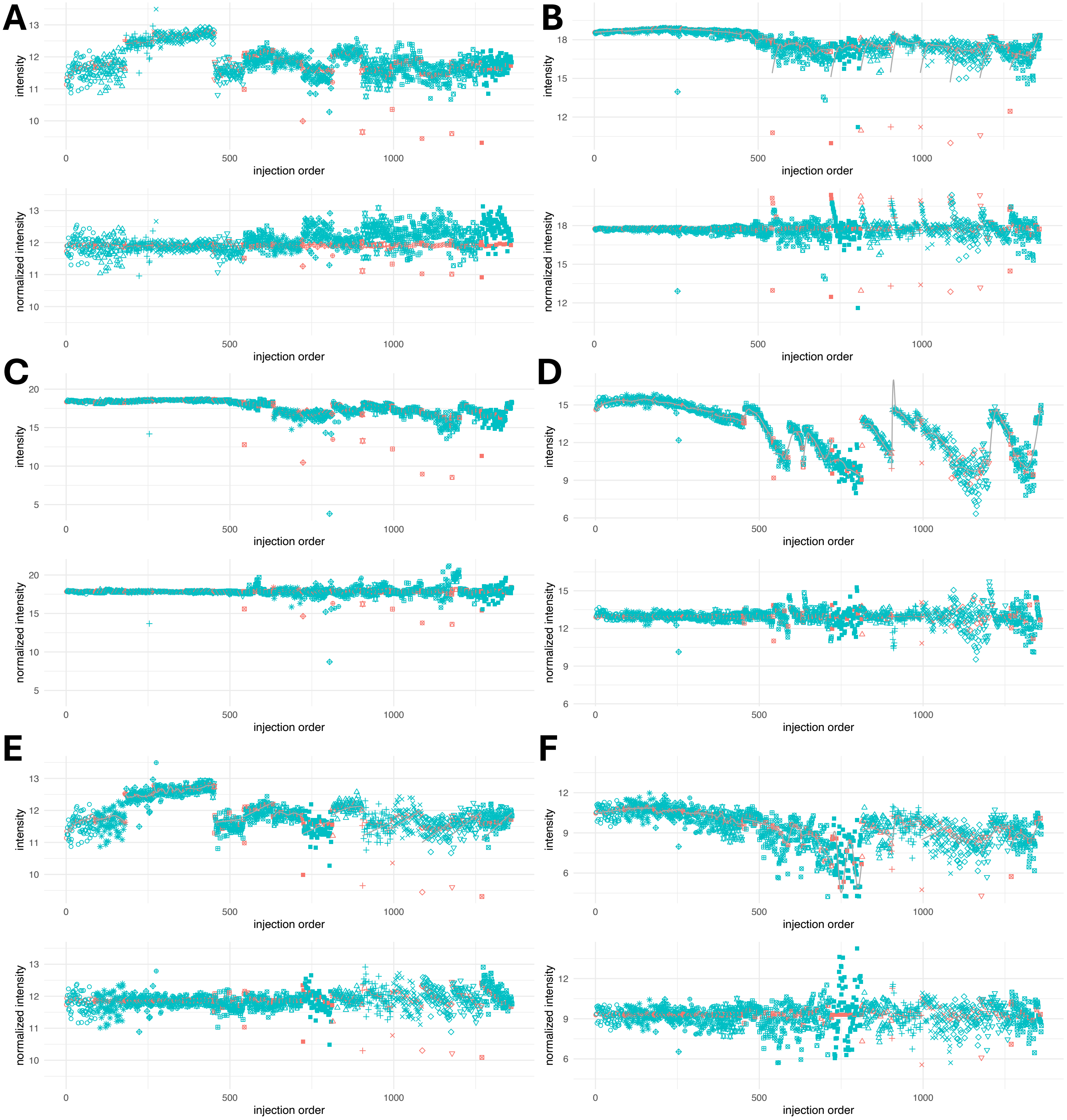


*Figure S8: Six cases of aberrant normalization (BioHEART dataset), each case showing data before (top panel) and after normalization (bottom panel). Blue symbols represent biological samples, coral-red symbols represent QC samples. Different symbols represent batches. (A) TIGER, with obvious remaining batch effects, (B) QC-RLSC with generalized cross-validation, showing the impact of outlying QC observations, (C) SERFF, with increased variance due to outlying observations (see also Fig. S5), (D) QC-RLSC with leave-one-out cross-validation, showing comparatively large sample deviations in proximity of sudden intensity changes, fitting with the methods' higher MSPE, (E) QC-RSC with generalized cross-validation, showing the impact of outlying QC observations, (F) QC-RSC with leave-one-out cross-validation, showing comparatively large sample deviations around injection 750, fitting with the methods' higher MSPE.*

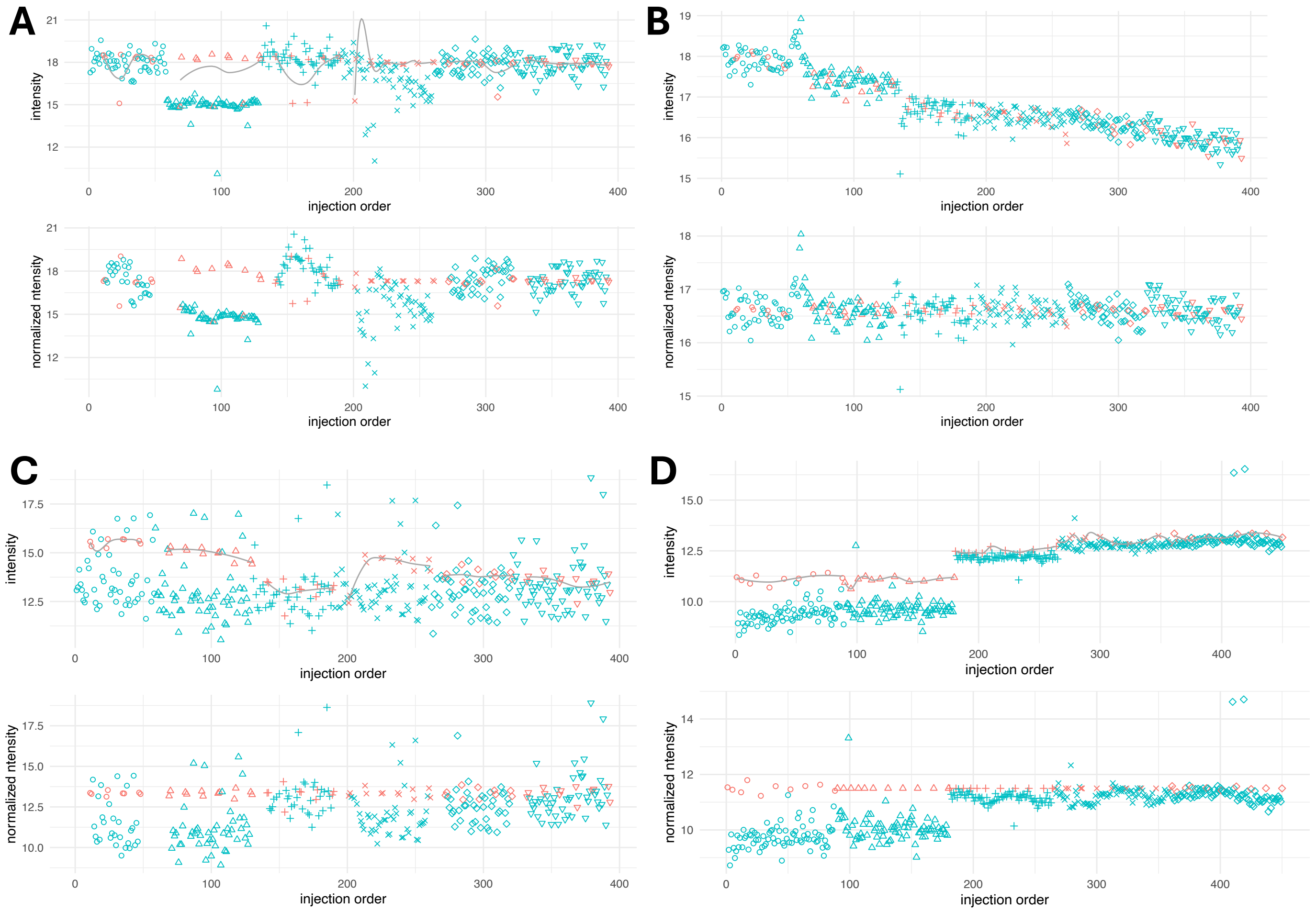


*Figure S9: Four examples of aberrant normalization. Blue symbols represent biological samples, coral-red symbols represent QC samples. Different symbols represent batches. (A) Raw intensities (top) and QC-RLSC normalized intensities (bottom) with leave-one-out cross-validation due to discrepancies in intensities observed between QC and non-QC runs, ENVIRONAGE dataset (B) Raw intensities (top) and SERFF-normalized intensities (bottom), showing remaining within-batch drift patterns, potentially due to an outlier value (see also Fig. S6 and S7), ENVIRONAGE dataset; (C) Raw intensities (top) and QC-RLSC normalized intensities (bottom) with leave-one-out cross-validation due to discrepancies in intensities observed between QC and non-QC runs, ENVIRONAGE dataset; (D) Raw intensities (top) and QC-RLSC normalized intensities (bottom) with leave-one-out cross-validation due to discrepancies in intensities observed between QC and non-QC runs, BioHEART dataset (first five batches).*

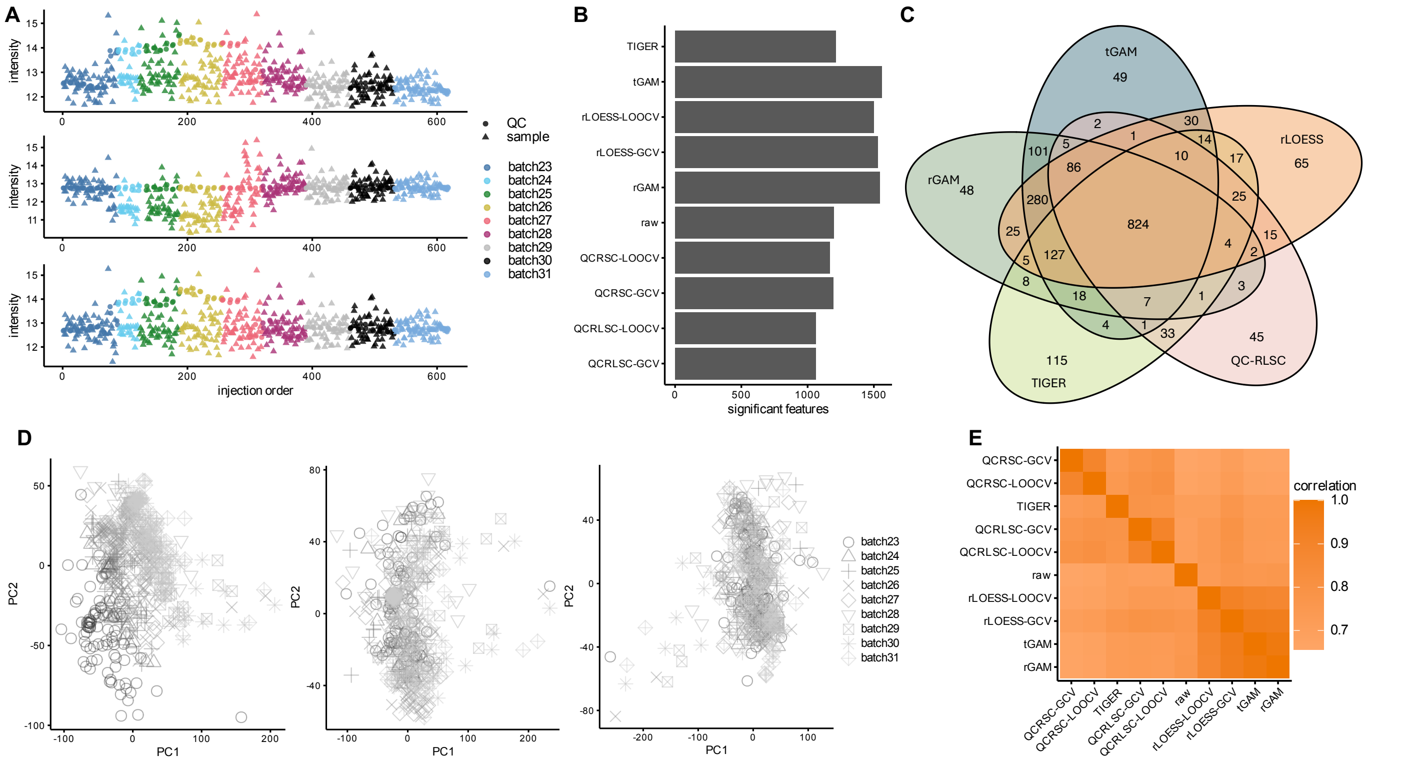


Figure S10: Impact of normalization methods in the FAME cohort. (A) Compound (ID3280) intensities before (top), after QC-RLSC (middle), and after tGAM (bottom) normalization. Note the discrepancies between QC and sample intensities in some batches, with QC intensities substantially above those in the samples, resulting in normalized sample intensities that are too low. (B) Number of significantly differentially abundant metabolites (DAMs) by normalization method. (C) Venn diagram of DAMs for the three robust methods, TIGER and QC-RLSC. (D) Principal component score plots before (left), after QC-RLSC (middle), and after tGAM (right) normalization. (E) Heatmap of correlations of p-values as obtained by Wilcoxon tests, per normalization method; methods ordered according to a hierarchical clustering.

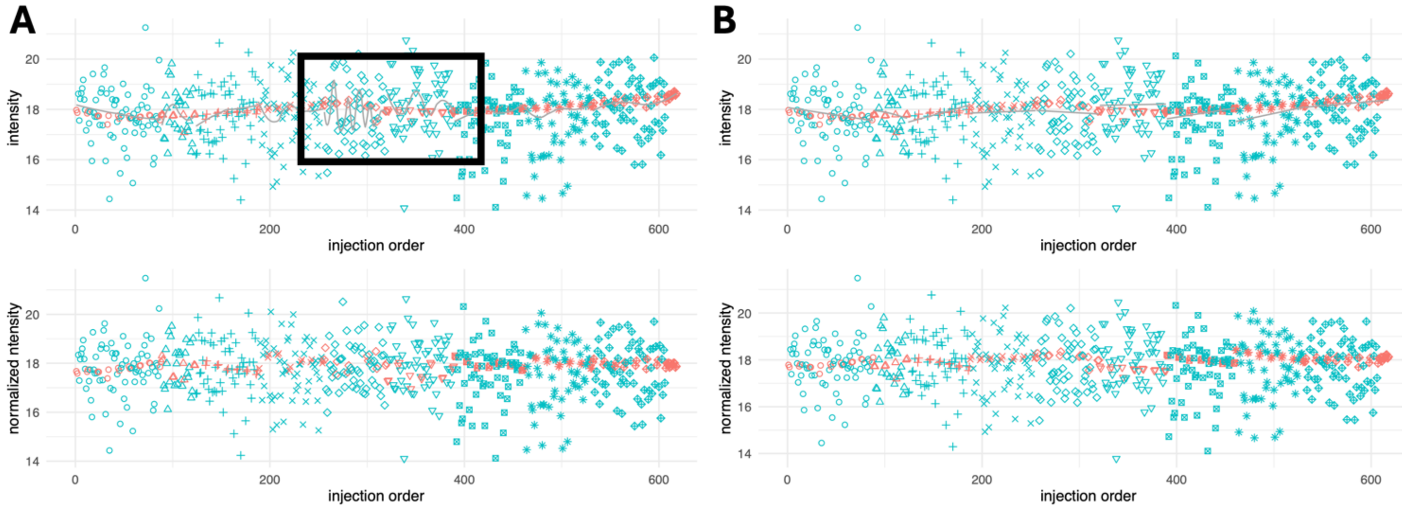


Figure S11: Metabolite ID86 in the FAME cohort, before (top panels) and after (bottom panels) normalization. Blue symbols represent biological samples, coral-red symbols represent QC samples. Different symbols represent batches. (A) rLOESS with generalized cross-validation normalization. Note the wiggly curve (boxed, black line). (B) tGAM normalization.

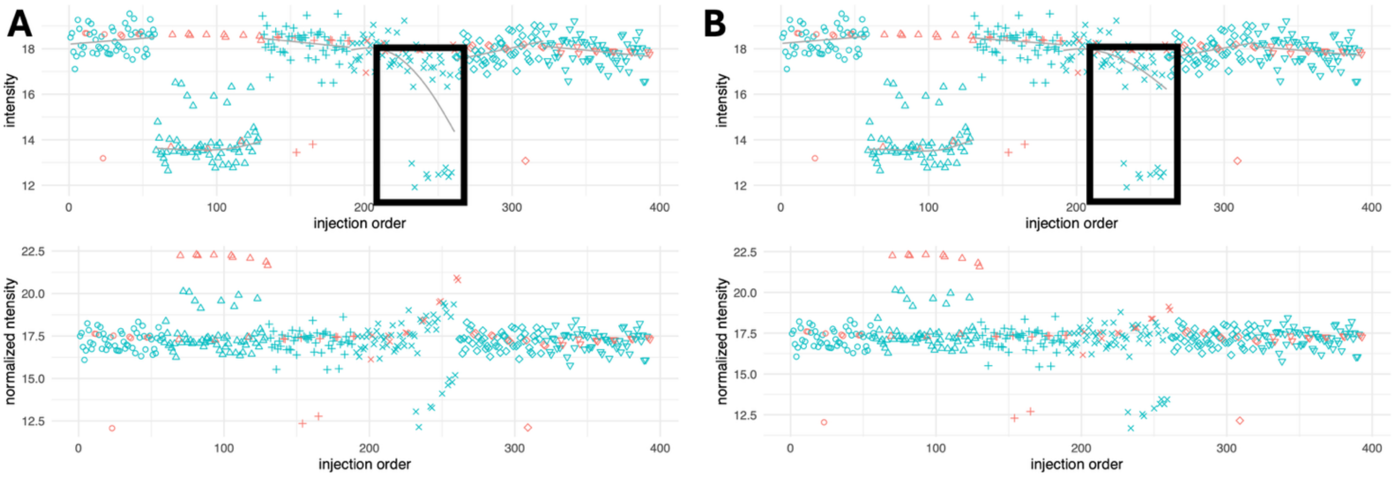


*Figure S12: Metabolite ID101 in the ENVIRONAGE cohort, before (top panels) and after (bottom panels) normalization. Blue symbols represent biological samples, coral-red symbols represent QC samples. Different symbols represent batches. (A) rGAM normalization, with a comparatively stronger impact of multiple observations with lower-than-typical intensities. (B) tGAM normalization, with a comparatively weaker impact of multiple observations with lower-than-typical intensities.*
